# Predicting interactions between the SARS-CoV-2 spike glycoprotein and the human proteome using AphaFold and molecular dynamics simulations

**DOI:** 10.1101/2025.07.28.667129

**Authors:** Max Henderson, Qun Liu

## Abstract

Since the COVID-19 pandemic began, the SARS-CoV-2 virus has caused over 775 million cases and more than 7 million deaths worldwide. Despite progress in treatments and vaccines, we still need better ways to prevent and treat the disease. To do this, a clearer understanding of how SARS-CoV-2 interacts with human proteins is needed. We developed a new computational tool to predict and study these protein-protein interactions. Our method uses two stages of AlphaFold, an AI tool, to predict how SARS-CoV-2 proteins bind to human proteins, followed by molecular dynamics simulation to refine these predictions. We tested this method in a small study focusing on the S1 subunit of the SARS-CoV-2 spike protein interacting with human cell junction and synaptic proteins, and in a larger study screening the spike S1 N-terminal domain against the entire human proteome. We validated the method using experimental virus–human protein interactions. The results provide valuable insights into how SARS-CoV-2 interacts with human proteins, guiding future experiments to better understand COVID-19’s short- and long-term effects. This developed method could be utilized to study protein-protein interactions in other biological systems.

## Introduction

The 2019 coronavirus disease (COVID-19) pandemic has impacted people from all over the world, with over 775 million cases and 7 million deaths globally [1]. In addition to the common physical symptoms of the disease, approximately 30% of those infected suffer from symptoms months and years after infection, which are deemed post-acute sequelae of COVID-19 (PASC) or “long-COVID”. Often these sequelae manifest as neurological symptoms such as fatigue, “brain fog”, sleep disturbances, difficulty concentrating, depression, and anxiety [2–6].

The causative agent of COVID-19 is severe acute respiratory syndrome coronavirus 2 (SARS-CoV-2). The viral genome encodes sixteen non-structural proteins, ten accessory proteins, and four structural proteins. These structural proteins consist of the spike glycoprotein (S), the membrane (M) protein, the envelope (E) protein, and the nucleocapsid (N) protein [7]. The glycosylated, homotrimeric spike protein, composed of the N-terminal S1 and C-terminal S2 subunits, is responsible for viral entry into cells. The S1 subunit is approximately 140 kDa and consists of an N-terminal domain (NTD) and a receptor-binding domain (RBD) that binds angiotensin converting enzyme 2 (ACE2) and is used for cellular entry [8]. The spike protein has thus been a protein of interest in combating COVID-19, leading to both full-length spike and RBD mRNA vaccines that have been developed and deployed [9]. In addition, antiviral therapeutics and monoclonal antibodies have been researched and developed that target the spike protein’s RBD binding to ACE2 [10]. Unfortunately, however, the spike protein, including the RBD, is prone to mutation, leading to immune escape and the genesis of new variants of concern [11–13]. Additionally, research suggests that another possible hurdle in the fight against COVID- 19 may be due to immune imprinting [14–19], perhaps due to spike and RBD mutations [20,21]. For these reasons, it is important to understand human–SARS–CoV–2 interactions and seek alternative therapeutic approaches to prevent SARS-CoV-2 infection as well as alleviate or cure COVID-19 and long-COVID symptoms.

In this study, we developed a computational workflow integrating AlphaFold [22] and computational techniques to explore the human–SARS–CoV–2 interactome to discover novel protein-protein interactions that may be involved in COVID-19 and long-COVID. AlphaFold was utilized to predict human–SARS–CoV–2 interactions [22]. Molecular dynamics (MD) simulations were then used to filter or refine the top AlphaFold predictions. Benchmarking was conducted to determine the workflow’s ability to narrow down many predictions into predictions that accurately match experimental protein-protein interactions. This computational tool could be used for predicting large-scale protein–protein interactions in the drug and target discovery toolkit.

## Results

### AlphaFold prediction of two known human-SARS-CoV-2 protein complexes

To test the validation of using AlphaFold2 for predicting SARS-CoV-2 interactions with human proteins, we performed predictions on the spike NTD with antibody 5-7 (PDB code 7RW2) and the E protein with PALS1 (PDB code 7M4R). The structures of both complexes were released in the PDB after the AlphaFold2 training date. We first used the full-length sequences for predictions, which did not predict desired interactions as shown in their complex structures.

When we truncated them into smaller regions, the results were dramatically improved. Using the amino acid sequence of antibody 5-7 that is involved in the interaction with the spike NTD, AlphaFold2 was able to correctly predict the interactions with a root mean square deviation (RMSD) of 0.763 Å when aligned to the crystal structure. The predicted aligned error (PAE) of the top prediction showed low PAE values, measured in angstroms, for the 5-7 helix with respect to both the hydrophobic cleft and the NTD as a whole (**Fig 1A**). Similarly, when making predictions using the E protein with no transmembrane region and a truncated human PALS1 (residues 248-336), AlphaFold predicted the correct orientation and position of the E interactions with PALS1 as shown in their cryoEM structure, with an RMSD of 0.838 Å. Accordingly, the AlphaFold prediction produced a low PAE, suggesting overall confidence in the relative position of each protein (**Fig 1B**). The two examples indicate the utility of AlphaFold in predicting SARS-CoV-2 human protein interactions.

**Fig 1.**
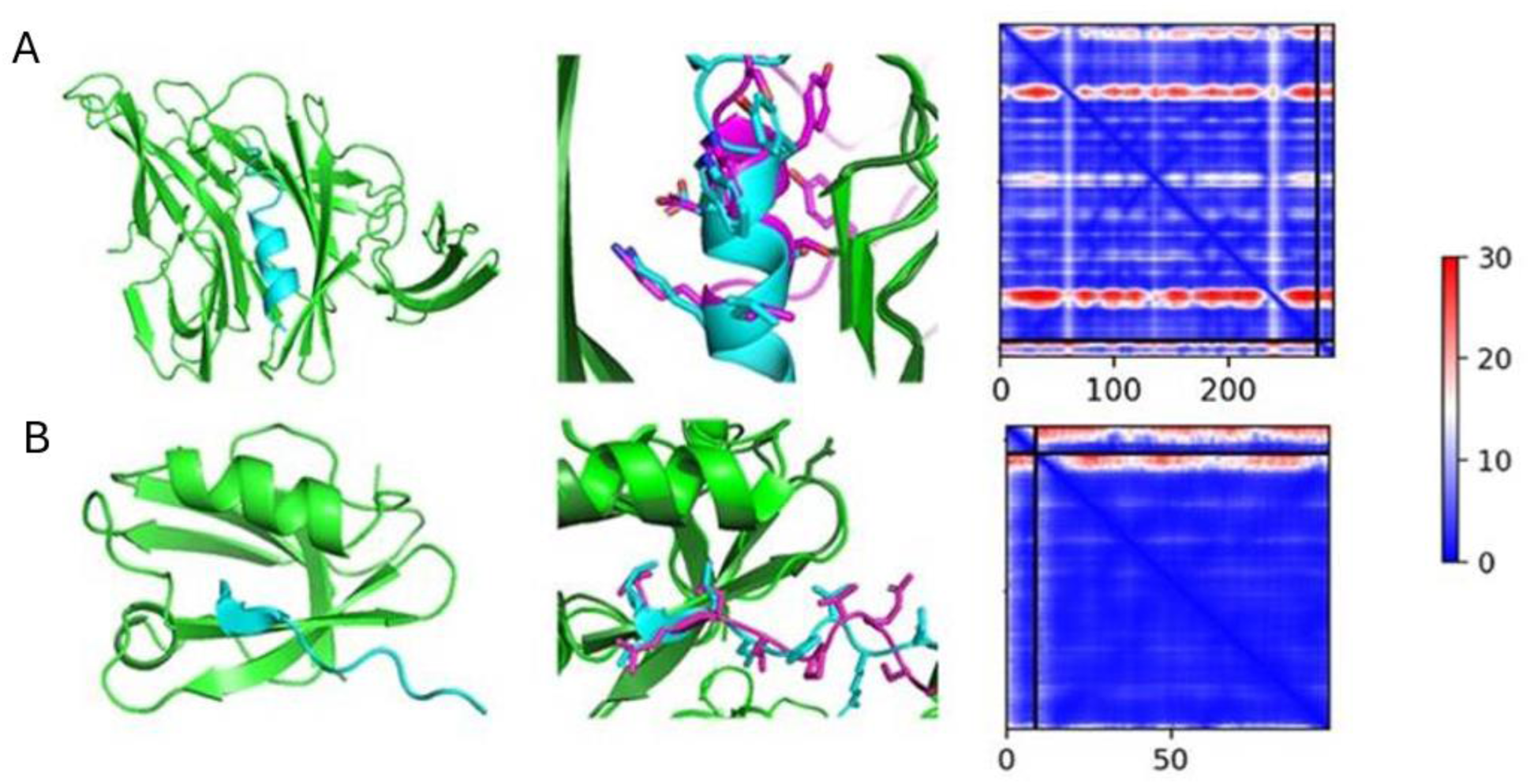
AlphaFold2 predictions of two controls suggest its ability to accurately predict SARS-CoV-2 and human interactions to some degree. A) left - predicted complex of antibody 5-7 (cyan) and NTD (green); middle – predicted complex aligned to experimental structure (7RW2; magenta, dark green); right – plot of PAE (large box – NTD; rectangle – antibody 5-7). B) left – predicted complex of E protein (cyan) and PALS1 (green); middle – predicted complex aligned to experimental structure (7M4R; magenta, dark green); right – plot of PAE (large box – PALS1; rectangle – E protein).

### AlphaFold prediction of additional known human-virus protein interactions

In order to determine the utility of AlphaFold in predicting protein–protein interactions between humans and viruses, we performed additional predictions on a set of 34 known structures of human-virus protein complexes that were released after the AlphaFold2 validation date (**Table S1**). This test set consisted of both SARS-CoV-2 and other viral structures. After the first round of predictions in which one unrelaxed model was created per heterodimer, an average RMSD of 14.79 ± 6.39 Å was obtained when alignments of the predicted models to their experimental structures. Additionally, the interaction surface areas were calculated for the experimental PDB structures to determine the range of empirical interaction surface areas between human and viral proteins. These interface area values range from 375.1 to 2887.5 Å^2^ with an average value of 951.2 Å^2^ (**Table S1**).

Based on initial results from the test set, we developed a quantitative filter using interface areas, clash scores, and pLDDT scores to identify high-confidence protein interaction models. For the interface area, we selected a threshold between the minimum and average values from the test set, ensuring moderate stringency. For the clash score, we set a cutoff of 20, as accurate predictions consistently had scores below this value. For pLDDT, although accurate predictions typically had scores above 80, we chose a cutoff of 70 to allow some flexibility while maintaining reliability. Applying these filters to all initial predictions, three models were selected for further analysis. 6YXJ is SARS-CoV macrodomain II in complex with human Paip 1 which aligned well with the experimental structure), 7RFY is importin alpha3 peptide in complex with MERS Orf4b which contained the wrong peptide orientation), and 7YUE is a single chain variable fragment targeted against a region of the SARS-CoV-2 Spike S2 subdomain which also had good alignment with its experimental structure. Additionally, three negative controls were selected. 7XBH is SARS-CoV-2 receptor binding domain bound to human ACE2, 7LMS is Coxsackievirus B3 2A protease bound to human SetD3, and 7X6Y is SARS-CoV-2 Nsp5 peptide bound to human TRIM7. 7XBH and 7LMS were selected due to their decent values and, when inspected visually, could look like accurate binding modalities, whereas 7X6Y was chosen due to its suboptimal filter values though the peptide was close to the correct binding pose.

We re-predicted the six complexes, generating five relaxed models per complex to increase structural diversity and reduce clash scores. All six complexes showed improved metrics after relaxation. The models for 6YXJ and 7YUE accurately predicted their structures, but 7RFY failed to predict the correct peptide orientation. Additionally, 7YUE produced a model with acceptable metrics but a flipped S2 fragment, which was used as an extra negative control.

Surprisingly, the negative control 7LMS achieved the correct binding pose; we selected both its correct pose (with the best metrics) and an incorrect model for further analysis. After molecular dynamics simulations, most models showed increased interaction surface areas, except the incorrect 7YUE model, which decreased from 824.8 to 624.7 Å² (**Table S2**). Although AlphaFold struggled to predict many test set interactions, our filtering criteria eliminated all but one poor prediction, yielding three accurate models (66% success rate).

We also analyzed how truncating protein sequences affects prediction accuracy, comparing PDB sequences (from experimental structures), full-length UniProt sequences, and custom truncations. Truncation improved predictions in some cases. For example, using the PDB sequence of 7DHG enabled AlphaFold to predict the correct binding orientation of Orf9, which was not achieved with the original PDB sequence, custom truncation, or full-length sequence. Similarly, the custom truncation of 7RF7 allowed an accurate prediction of Orf4b’s binding pose, missed by other sequence versions (**Table S3**).

### Empirical computational workflow to predict SARS-CoV-2 spike – human protein interactions

Based on the AlphaFold prediction of known viral and human protein-protein interactions, we proposed a computational workflow to predict SARS-CoV-2 spike interactions with human proteins (**Fig 2**). This workflow begins with an initial set of predictions using a protein of interest against a collection of potential interactors. This initial screen will produce one unrelaxed model per heterodimer in order to reduce the computational cost. These initial predictions are then analyzed and filtered by interaction surface area, clash score, and pLDDT score. Those predictions that make it through this quantitative filtering are then visually inspected to determine whether the predicted structure appears realistic. Next, the sequences of the selected predictions are then predicted again for AlphaFold2 prediction, allowing for five relaxed models. The filtering and analysis are then performed on all five predicted models for each heterodimer. Molecular dynamics simulations are then performed on the selected predictions to refine and optimize the interactions.

**Fig 2.**
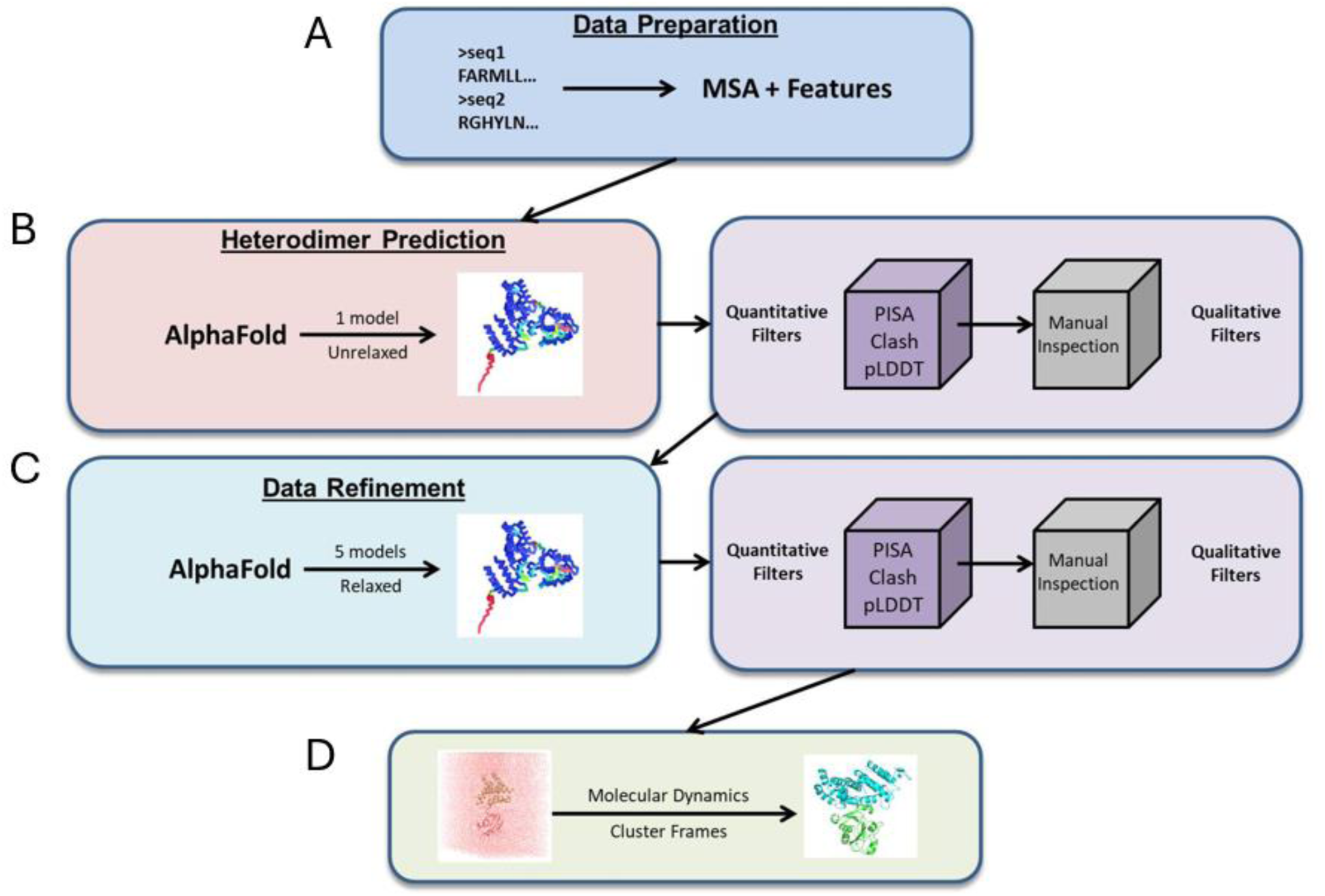
Computational workflow to predict protein-protein interactions. The workflow consists of four steps: A) Data preparation. B) Initial interaction predictions and analysis and filtering of the interactions through quantitative and visual filters. Interaction surface area, intermolecular clash, and pLDDT scores, visual inspection, and functional annotation. C) Data refinement and refiltering. D) Molecular dynamics simulation. Molecular dynamics simulations to obtain physics-aware models of the predicted interactions.

### SARS-CoV-2 Spike S1 interactions with human cell junction and synaptic proteins

In the initial phase of Spike S1 screening, we generated one unrelaxed heterodimeric model for each of approximately 2,700 human cell junction and synaptic protein sequences, reducing to about 2,500 sequences due to prediction limitations from sequence length constraints and the presence of noncanonical amino acids. Subsequent filtering using three criteria further reduced the number of predictions. First, we retained models with interface areas greater than 500 Å², narrowing the predictions to approximately 1,100 [23,24]. Next, applying a Phenix clash score cutoff of 20 reduced these to about 200 unrelaxed models [25]. We then visually inspected these 200 models, discarding those that appeared structurally unrealistic or lacked relevant biological functions based on UniProt annotations, resulting in 20 models. These 20 human proteins were conservatively truncated to their apparent interacting regions and re-predicted against Spike S1 using AlphaFold2. The seven models with the largest interface areas and lowest clash scores were designated as “hits” (**Fig. 3A-G**, **Table 1**).

**Fig 3.**
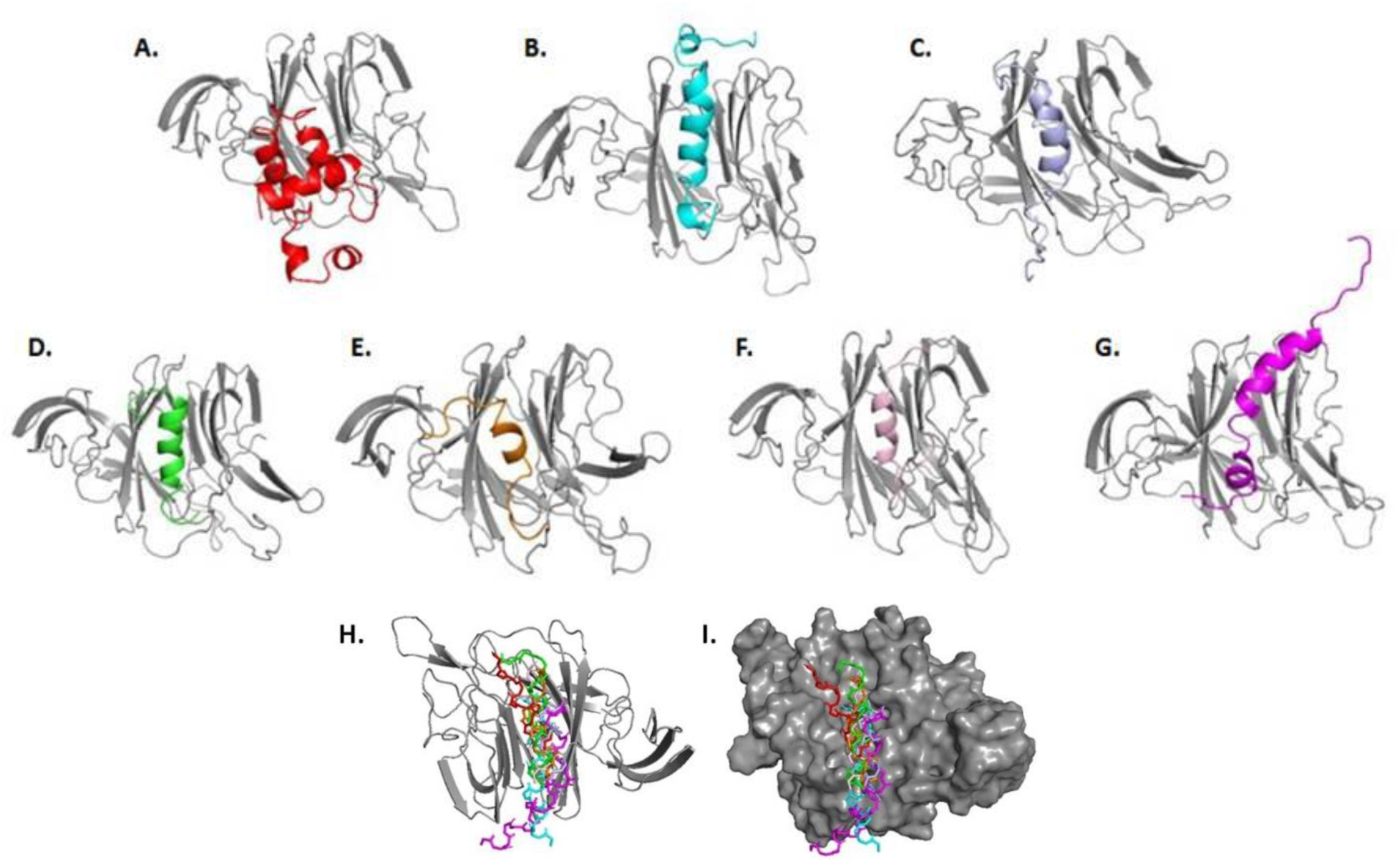
**Seven human proteins predicted to bind to a common cleft in the spike N-term domain**. Representative structures from molecular dynamics simulations with NTD are shown in grey. A) α-neoendorphin (red); B) α-synuclein (cyan); C) synaptotagmin-1 (pale blue); D) neuropeptide K (green); E) paxillin (orange); F) presenilin-1 (pink); G) BNIP3 (purple). Panels H and I show the superimposition of the backbone of the interacting regions of A – G, with NTD displayed as both cartoon (H) and surface (I) views.

**Table 1.**
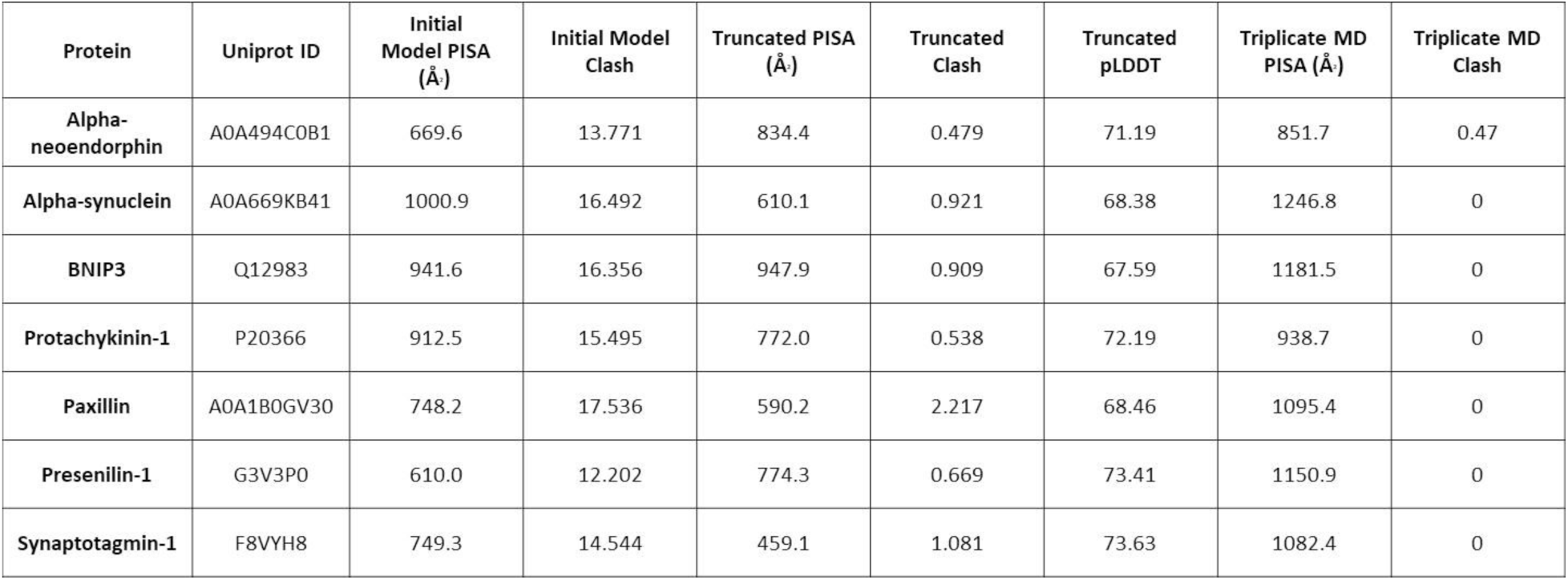
Interaction surface area and clash scores for the seven hits from the human cell junction and synaptic proteins.

| Protein | Uniprot ID | Initial Model PISA (Å) | Initial Model Clash | Truncated PISA (Å) | Truncated Clash | Truncated pLDDT | Triplicate MD PISA (Å) | Triplicate MD Clash |
| --- | --- | --- | --- | --- | --- | --- | --- | --- |
| Alpha-neoendorphin | A0A494C0B1 | 669.6 | 13.771 | 834.4 | 0.479 | 71.19 | 851.7 | 0.47 |
| Alpha-synuclein | A0A669KB41 | 1000.9 | 16.492 | 610.1 | 0.921 | 68.38 | 1246.8 | 0 |
| BNIP3 | Q12983 | 941.6 | 16.356 | 947.9 | 0.909 | 67.59 | 1181.5 | 0 |
| Protachykinin-1 | P20366 | 912.5 | 15.495 | 772.0 | 0.538 | 72.19 | 938.7 | 0 |
| Paxillin | A0A1B0GV30 | 748.2 | 17.536 | 590.2 | 2.217 | 68.46 | 1095.4 | 0 |
| Presenilin-1 | G3V3P0 | 610.0 | 12.202 | 774.3 | 0.669 | 73.41 | 1150.9 | 0 |
| Synaptotagmin-1 | F8VYH8 | 749.3 | 14.544 | 459.1 | 1.081 | 73.63 | 1082.4 | 0 |

### Predicted cell junction and synaptic proteins bind to a conserved cleft in the N-terminal domain

Interestingly, all seven hits interact with the same region of the S1’s N-terminal domain (NTD). After aligning the sequences of the NTD of SARS-CoV-2 variants, we found that the NTD is highly conserved (**Fig S1**). This interaction site is a hydrophobic cleft (**Fig S2**). Additionally, all hits interact with the cleft via their helical regions (**Fig 3H and I**).

The prediction of the seven hits using their truncated regions resulted in increased (α- neoendorphin, BNIP3, and Presenilin-1) and decreased (α-synuclein, Protachykinin-1, Paxillin, and Synaptotagmin-1) interface areas. Additionally, these predictions using truncation had reduced clash scores ranging from 0.5 to 2 (**Tables 1 and S4**).

The predicted NTD models normally have higher pLDDT scores (79-82) compared to those of human proteins (30-57) [26]. This discrepancy may be expected as some of these hits do not have experimental structures or are disordered, such as neuropeptide K [27], leading to lower pLDDT scores [28]. Interestingly, however, PAEs of the top models have low PAEs for the interacting regions (**Fig S4**), suggesting a decent quality of our predictions.

MD simulations were then performed on the seven models in triplicate. We aligned the representative structures of the triplicates (**Fig 3A-G**) with the AlphaFold2 models, and we found that both interface areas and clash scores were improved. For example, the AlphFold2 models have interface areas ranging from approximately 450 to 950 Å^2^. After MD simulations, interface areas increased to a range of 850 to 1250 Å^2^ (**Table 1**).

We analyzed the N-terminal domain (NTD) interaction regions for the seven identified hits. The interaction region of the paxillin isoform (residues 44–58) corresponds to residues 262–276 of the canonical paxillin (PAXI, UniProt ID: P49023) and includes the LD4 motif. Residues 23–41 of the α-neoendorphin isoform match the propeptide residues 23–41 of canonical α- neoendorphin (PDYN, UniProt ID: P01213). The synaptotagmin-1 isoform residues 40–50 align with luminal residues 40–50 of canonical synaptotagmin-1 (SYT1, UniProt ID: P21579).

Protachykinin-1 (TAC1) residues 77–88 are part of the neuropeptide K (NPK, residues 72–107) cleavage product. Residues 8–18 of the presenilin-1 isoform correspond to cytoplasmic residues 8–18 of canonical presenilin-1 (PSEN1, UniProt ID: P49768). Finally, residues 82–94 of the α-synuclein isoform match residues 82–94 of canonical α-synuclein (SNCA, isoform-1, UniProt ID: P37840-1).

### Large-scale prediction of Spike NTD interactions with the human proteome

Building on promising results from small-scale screening, we expanded N-terminal domain (NTD) interaction predictions to the human proteome, starting with approximately 20,500 protein sequences. We filtered out sequences longer than 2,500 residues, those with noncanonical amino acids, and those affected by computational errors, yielding about 19,700 initial predictions. To ensure high-confidence results, we applied strict filters: interface areas greater than 500 Å², Phenix clash scores below 10, and pLDDT scores above 80, supplemented by biological annotations [26]. Selected hits were re-predicted using AlphaFold2, generating five relaxed models per prediction [29].

From the large-scale predictions, 330 unrelaxed models met the filtering criteria. Of these, 126 had reasonable structures, and 58 had relevant functional annotations. Re-prediction of these 58 sequences with AlphaFold2 produced 24 models that passed the filters. Visual inspection identified 17 models with at least one structurally realistic structure. These final hits included five T cell receptor variable regions, three interferon alphas, one leukocyte immunoglobulin-like receptor, two prostate and testis-expressed proteins, one translocon-associated protein subunit, one trefoil factor, one tetratricopeptide repeat protein, one exopolyphosphatase, and one sorting nexin-6 (**Table S5**).

Upon a review of the literature, functionally, five T cell receptor regions and three interferon alphas appear to play some roles in the immune response to COVID-19 (T cell receptor regions [30–34]; interferons [35–37]). Additionally, a BIOGRID search showed evidence of spike interactions with sorting nexin-6 and the translocon-associated protein subunit [38]. The BIOGRID search also showed that the spike protein interacts with other tetratricopeptide repeat proteins, though the protein predicted here was not listed. Thus, it appears the literature provides support for predicted human proteins involved in COVID-19.

The predicted NTD interactions involve two binding sites: the cleft and an N-terminal β-sheet (**Fig. S5A-B**). To eliminate hits clashing with the spike trimer, we aligned the predicted models with experimental spike structures, retaining eight clash-free hits that avoided neighboring spike domains or glycans. Hits interacting with the β-sheet, primarily T cell receptors, could not be accommodated due to spatial constraints, suggesting possible S1 shedding for these complexes. Of the eight hits, four (interferon alphas and prostate and testis-expressed protein 4) had high-quality PAE values, three (tetratricopeptide repeat protein, trefoil factor 2, and testis-expressed protein 3) had medium PAE values, and one (PRUNE1) had poor PAE values (**Fig. S5C-F**). MD simulations of the seven high- and medium-quality hits showed varied interface area changes (**Fig. 4**, **Table 2**). Interface areas increased for interferon alpha-14 (862 to 966.6 Å²), tetratricopeptide repeat protein 39A (632 to 1730.5 Å²), and trefoil factor 2 (738.9 to 1044.1 Å²). They slightly decreased for interferon alpha-1/13 (885.3 to 761.3 Å²), interferon alpha-2 (963.9 to 827.8 Å²), and prostate and testis-expressed protein 4 (972.8 to 795.1 Å²), while decreasing significantly for testis-expressed protein 3 (608.9 to 278.2 Å²).

**Fig 4.**
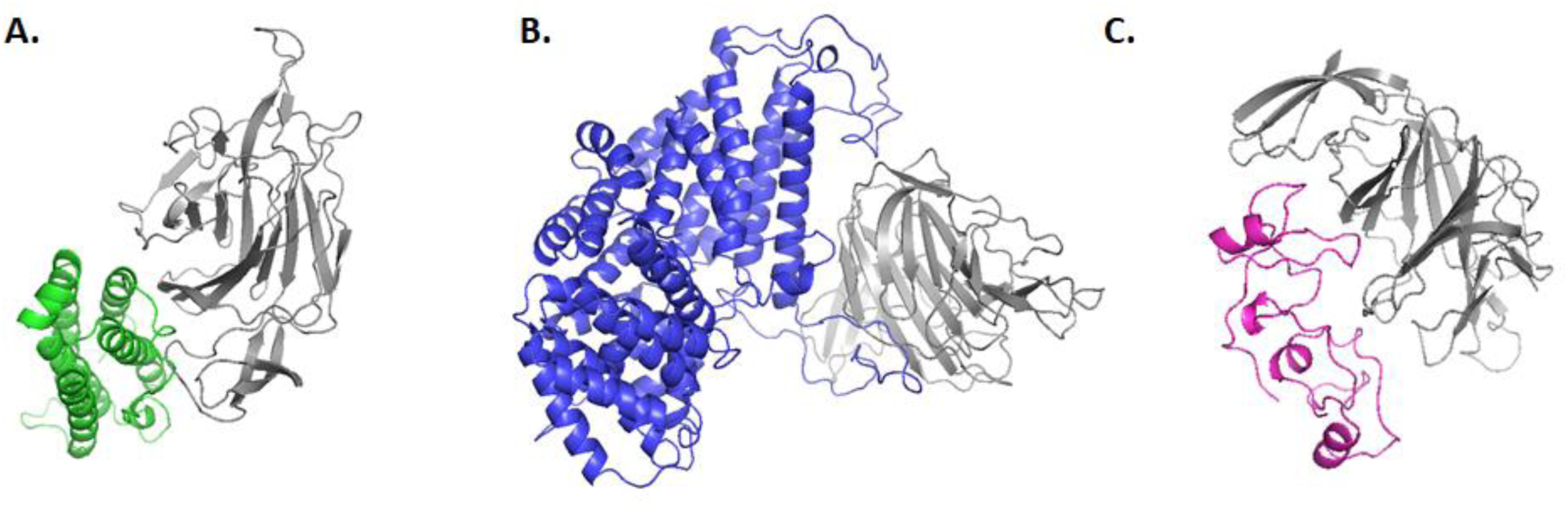
**Interactions between spike NTD and human proteins**. Depicted are representative structures of A) interferon alpha-14 (Uniprot# P01570, green); B) tetratricopeptide repeat protein 39A (Uniprot# Q5SRH9, blue); C) trefoil factor 2 (Uniprot# Q03403, magenta). Spike NTD is shown as gray cartoons.

**Table 2.** Comparison of interface areas and clashes before and after molecular dynamics simulation for the large-scale screening hits. Interface area improved for P01570, Q5SRH9, and Q03403, slightly decreased for P01562, P01563, and P0C8F1, and dramatically decreased for B3GLJ2.

| Name | Uniprot ID | Best Visual Model | PISA (Å<sup>2</sup>) | Total Clash | MD PISA (Å<sup>2</sup>) | MD Total Clash |
| --- | --- | --- | --- | --- | --- | --- |
| Interferon alpha-1/13 | P01562 | rank_1_model_3 | 885.3 | 0.886 | 761.3 | 0 |
| Interferon alpha-14 | P01570 | rank_1_model_3 | 862 | 0.432 | 966.6 | 0.441 |
| Interferon alpha-2 | P01563 | rank_1_model_3 | 963.9 | 0.411 | 827.8 | 0 |
| Prostate and testis expressed protein 3 | B3GLJ2 | rank_3_model_4 | 608.9 | 0 | 278.2 | 0 |
| Prostate and testis expressed protein 4 | P0C8F1 | rank_1_model_4 | 972.8 | 0.819 | 795.1 | 0 |
| Tetratricopeptide repeat protein 39A | Q5SRH9 | rank_2_model_5 | 632.1 | 0.427 | 1730.5 | 0 |
| Trefoil factor 2 | Q03403 | rank_3_model_3 | 738.9 | 1.023 | 1044.1 | 0 |

## Discussion

COVID-19 is still a global concern. Recent variants of concern or interest, such as JN.1, are still appearing and spreading [1,39]. While therapeutics that focus on the receptor-binding domain of the spike glycoprotein have been a large focus, there is a need for a broader range of drug targets due to the RBD being prone to mutation and immune escape [11–13]. To address this, we have created a computational workflow and used it to predict novel protein–protein interactions between spike and human proteome in hopes of adding to the list of potential druggable targets (**Fig 2**).

This work consisted of two primary studies: a small-scale study focusing on the spike S1 subunit and human cell junction and synaptic proteins, as well as a large-scale study where the spike N-terminal domain (NTD) and the human proteome were used. In the first study, the application of the workflow on the approximately 3000 synaptic and cell junction proteins filtered down the number of proteins to seven potential hits. Interestingly, all of the human proteins in the small-scale study were predicted to interact with a hydrophobic cleft of the spike S1’s NTD (**Fig 3**).

While the overall predictions had modest pLDDT scores, that may be expected because some hits appear to be unstructured in solutions. For example, neuropeptide K has no clear secondary structure until it is in the presence of micelles [27]. Additionally, examining PAEs shows a much smaller error for the interacting residues (**Fig S4**). Further supporting that some of these hits may interact with this hydrophobic cleft are the experimental structures that have been solved of heavy complementarity-determining region 3 loop of monoclonal antibody PVI.V6-14 Fab (7RBU, [40]), polysorbate 80 (7JJI, [41]), biliverdin (7B62, [42]), and the heavy complementarity-determining region H3 of neutralizing antibody 5-7 Fab (7RW2, [43]).

Interestingly, it appears some of these molecules show the ability to bind and neutralize multiple SARS-CoV-2 variants, likely due to the NTD being highly conserved (**Fig S1 and S6**) [40,43]. The results from the small-scale study may provide new avenues for research in long-COVID as there has been possible viral detection in the central nervous system of both humans and non-human primates [44–49]; for reviews see [50–52]. The S1 subunit may impact brain endothelial cells, such as in the blood-brain barrier (BBB) [53–56]. Additional research has shown that the S1 subunit can also cross the BBB in mice with the entry of the S1 subunit into mouse and rat brains leading to behavioral changes, cognitive deficits, and negative structural and biological changes [57–60].

In our large-scale study, we screened approximately 20,000 human proteome proteins, narrowing them down to 17 hits (Table S5). These hits included T cell receptor variable regions, interferon alphas, a leukocyte immunoglobulin-like receptor, prostate and testis-expressed proteins, a translocon-associated protein subunit, a trefoil factor, a tetratricopeptide repeat protein, an exopolyphosphatase, and sorting nexin-6. Molecular dynamics simulations further refined this list.

Notably, AlphaFold2 struggled to predict the interaction between the SARS-CoV-2 Spike protein (specifically the receptor-binding domain) and human ACE2, possibly due to heavy glycosylation on both proteins, with some glycans likely influencing binding [61–64].

Nevertheless, this study identifies 14 novel targets for COVID-19 research. While our benchmarking validated the workflow’s ability to predict human–virus protein interactions, these predictions require experimental validation. This workflow could be adapted for broader biological systems, serving as a valuable tool for studying protein–protein interactions in various research contexts.

## Methods

### Sequence Collection

The Delta variant (B.1.1.529) spike protein sequence was obtained from GenBank (UFO69279.1). The S1 subunit (residues 14-695) of the spike protein was obtained by truncating the protein after the furin cleavage site and removing the signal sequence. The human protein sequences were obtained from UniProt using the search terms “synapse” and “junction.” The human synapse (SL-0258), human paranodal septate junction (SL-0192), and human cell junction (SL-0038) libraries, including isoforms, were downloaded in FASTA format. Finally, the individual FASTA files were combined into a single file (2820 total sequences at the time of collection). For the Summit work, the FASTA files for the human proteome (UP000005640) were retrieved from UniProt with one protein per gene, leading to a total of 20,598 protein sequences. All sequences were retrieved between the years 2021 and 2022.

### Experimentally determined SARS-CoV-2 – human protein interactions

Sequences of the SARS-CoV-2 spike protein (P0DTC1), envelope protein (P0DTC4), and human PALS1 (Q8N3R2) were obtained from Uniprot. The antibody 5-7 sequence was taken from the spike – antibody 5-7 complex structure (PDB code: 7RW2). Predictions were performed using a local version of ColabFold [29].

### Benchmarking Dataset

The benchmarking test dataset structures were accessed between the dates of Feb 7, 2024 and Feb 12, 2024 by searching the PDB with criteria including combinations of the structural keywords of “viral protein” and/or source organism of “homo sapiens,” “10239,” and/or “2501931.” All searches were limited to distinct protein entities = 2. The PDB entry release date was set for after AlphaFold2’s validation date of Feb 15, 2021. Duplicates as well as chimeric entities, capsids, and dimeric antibodies were all removed. Additionally, SARS-CoV-2 spike– human ACE2 structures were overrepresented in the initial set, so many of these were removed. By the end of test dataset curation, we obtained a total of 35 PDBs of virus–human protein interactions, including 18 SARS-CoV-2 structures (13 spike-related and 5 containing non-spike structures) and 19 non-SARS-CoV-2 viral proteins.

Benchmarking was performed on the test dataset using the same workflow as described in this paper. At each stage of the workflow, RMSD was calculated between the predicted and experimental structures using PyMOL [65].

### Predict Protein-protein interactions

Multiple sequence alignment (MSA) and structure prediction were performed using a locally installed version of ColabFold’s AlphaFold2_advanced notebook [29]. MSA was performed using MMSeqs2 on the truncated spike S1 sequence in combination with the combined FASTA library sequences described above [66]. The minimum sequence length was 16 and the maximum sequence length was 2500 residues for MSA. No templates were used for structure prediction.

The oligomeric state was set as 1:1. The ColabFold scripts were modified to allow it to run on GPU clusters. To reduce the computational time, the initial heterodimer predictions were set to output one model without relaxation. For the Summit work, MSAs were precomputed locally and predictions were made using Summit’s AlphaFold-multimer container, which leads to one unrelaxed model per prediction. Predictions were made using the spike NTD (14-275).

### Analysis of predicted protein-protein interactions

The surface area of the protein-protein interactions was calculated using the CCP4 program PISA [23,24]. Additionally, the ‘clash score’ of each prediction was calculated using the Phenix software package [25]. The clash score is calculated first by determining the total number of nonbonded contacts of atomic radii with overlaps greater than 0.4Å, normalizing this value over the total number of atoms in the protein, and finally multiplying by 1000, meaning the clash score represents total clashes per 1000 atoms (1000 * (total number of bad overlaps/total number of atoms)). The intermolecular clashes between the spike and human proteins were extracted and the total scores were recalculated. Finally, the predictions were sorted using PISA and clash score cutoffs of over 500 Å^2^ and under 20 Å, respectively. The selected predictions after applying the cutoff were then visually inspected. The biological relevance of human proteins was then obtained using their UniProt annotations.

### Optimize protein-protein interactions

After the prediction and selection of spike-human synaptic/cell junction interactions, the seven hits were truncated to focus on their interacting regions. We removed the transmembrane domains and large non-interacting flexible regions. Additionally, truncations to the test set showed the potential for teasing out previously unpredicted, accurate models. Specifically, BNIP3 (Q12983; 194 aa) was truncated to residues 14-126, which included the BH3 domain (100-125). Alpha-neoendorphin (fragment A0A494C0B1; 89 aa) was truncated to residues 1-71, a region included in the canonical (P01213) propeptide region. Alpha-synuclein (isoform A0A669KB41; 151 aa) was truncated to residues 1-100. Paxillin (isoform A0A1B0GV30; 207 aa) was truncated to residues 40-60, which includes LD Motif 4 (residues 265-276 of canonical P49032). Presenilin-1 (fragment G3V3P0; 62 aa) was truncated to residues 1-25, part of the cytoplasmic region of the canonical form (residues 1-82 of P49768). Protachkynin-1 (P20366; 129 aa) was truncated to residues 72-107, which includes the sequence of Neuropeptide K (72- 107). Finally, Synaptotagmin-1 (fragment F8VYH8; 58 aa) was truncated to residues 31-58, part of the vesicular domain of the canonical form (residues 1-57 of P21579). These truncated protein sequences were then subject to the prediction workflow, with the output being five relaxed models. Alpha-neoendorphin, paxillin, presenilin-1, protachykinin-1, and synaptotagmin-1 had no further truncations. The truncated human proteins were then run through the same heterodimer prediction process as described above, except that each prediction produced 5 relaxed models ranked by per-residue confidence (pLDDT) score [26].

### Molecular dynamics simulations

For molecular dynamics, BNIP3 was truncated further to residues 91-126 and alpha-synuclein was truncated to residues 66-100. The selected Summit predictions were not truncated before molecular dynamics simulations. Molecular dynamics simulations were conducted using Schrodinger’s Desmond [67]. For simulations, the Delta variant spike NTD (14-275) was used (B.1.1.529; UFO69279.1). This was considered reasonable as various SARS-CoV-2 spike NTDs of different lengths have been stably expressed and purified [42,43,68,69]. The system for all simulations used the SPC water model and the OPLS_2005 force field in an orthorhombic solvent box. Each of the benchmarking and small-scale simulations was run in triplicate, containing 150 mM NaCl as well as neutralizing ions, and had a simulation time of 100 nanoseconds. The human proteome molecular dynamics were set up in the same way, with a single run of 100 nanoseconds.

### Analysis of molecular dynamics trajectories

Representative structures for each simulation of the small-scale study, as well as the benchmarking study, were obtained using Desmond’s ‘Desmond Trajectory Clustering’ task. First, the three replicas of each simulation were merged into one trajectory and aligned to the first frame, leading to a total of 3000 frames. For clustering, the RMSD Matrix was calculated using the backbone atoms of each truncated hit with a frequency of 30 frames. The most populated cluster was considered the representative structure. The process was also used for the Summit results by clustering 1000 frames with a frequency of 10.

## Acknowledgements

We thank Dima Kozakov and Carlos Simmerling for helpful suggestions and Michael Sandoval for assistance with the Summit work using the OCLF at ORNL.

## Author contributions

M. H. and Q. L. designed the experiment, performed the experiments, analyzed the data, and wrote the manuscript.

## Data Availability

All AlphaFold predictions are available upon reasonable requests.

## Supplemental Figures and Tables

**Fig S1.**
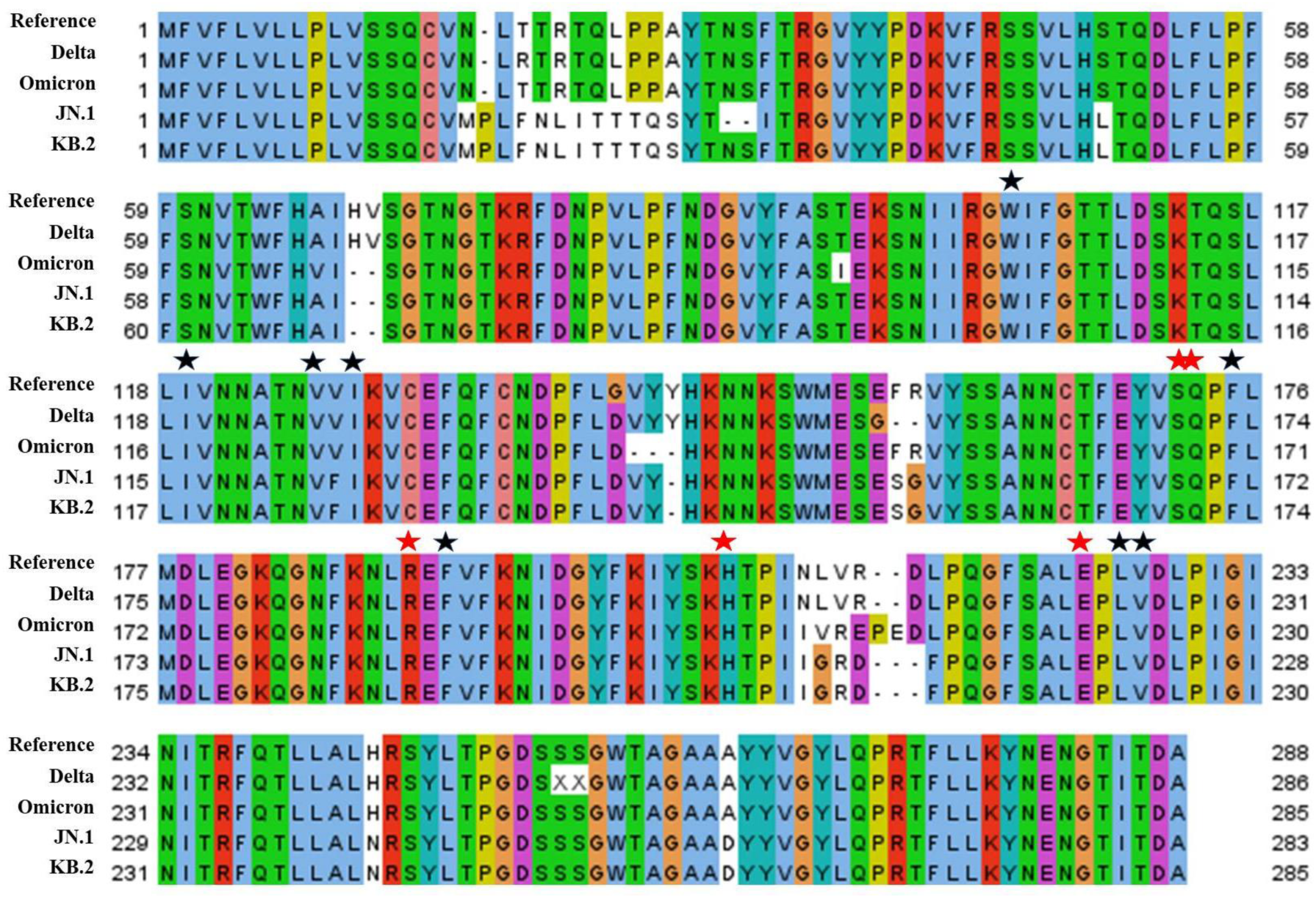
**Sequence alignment of multiple SARS-CoV-2 variants’ spike N-terminal domains suggest the domain is highly conserved**. 100% sequence identity shown in dark blue. Black and red stars indicate hydrophobic and electrostatic contacts, respectively, with at least one experimental structure (7RBU, 7JJI, 7B62, 7RW2) and one or more of the seven hits. 2019 Reference (NCBI: YP_009724390.1; REFSEQ: NC_045512.2); B.1.617.2 Delta (GenBank: PQ169688.1; Protein ID: XDZ89950.1); BA.1 Omicron (GenBank: PQ173214.1; Protein ID: XEA31857.1); JN.1 (GenBank: PQ168624.1; Protein ID: XDZ85688.1); KB.2 (GenBank: PQ199741.1; Protein ID: XEM76582.1).

**Fig S2.**
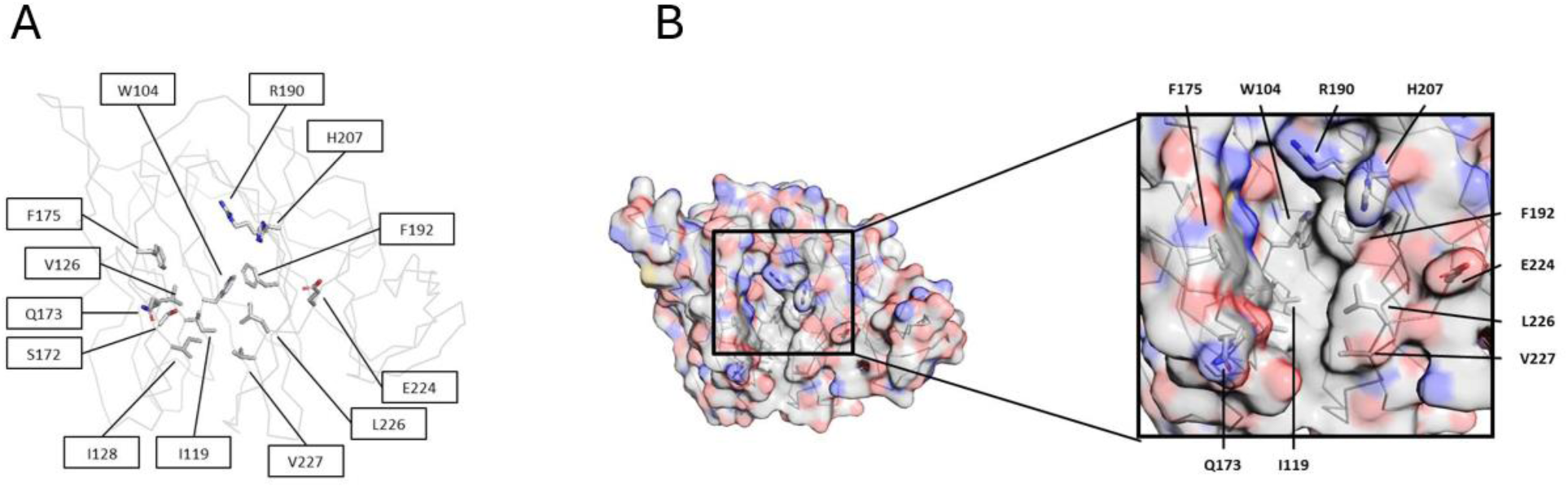
**The predicted binding site of the seven hits is a hydrophobic pocket surrounded by a few electrostatic regions**. A) Ribbon view of 7RW2 NTD with side chains of interacting residues shown (from Fig S1). B) Displayed is the electrostatic surface of the spike NTD from PDB 7RW2. Neutral (white); Positive (blue); Negative (red).

**Fig S3.**
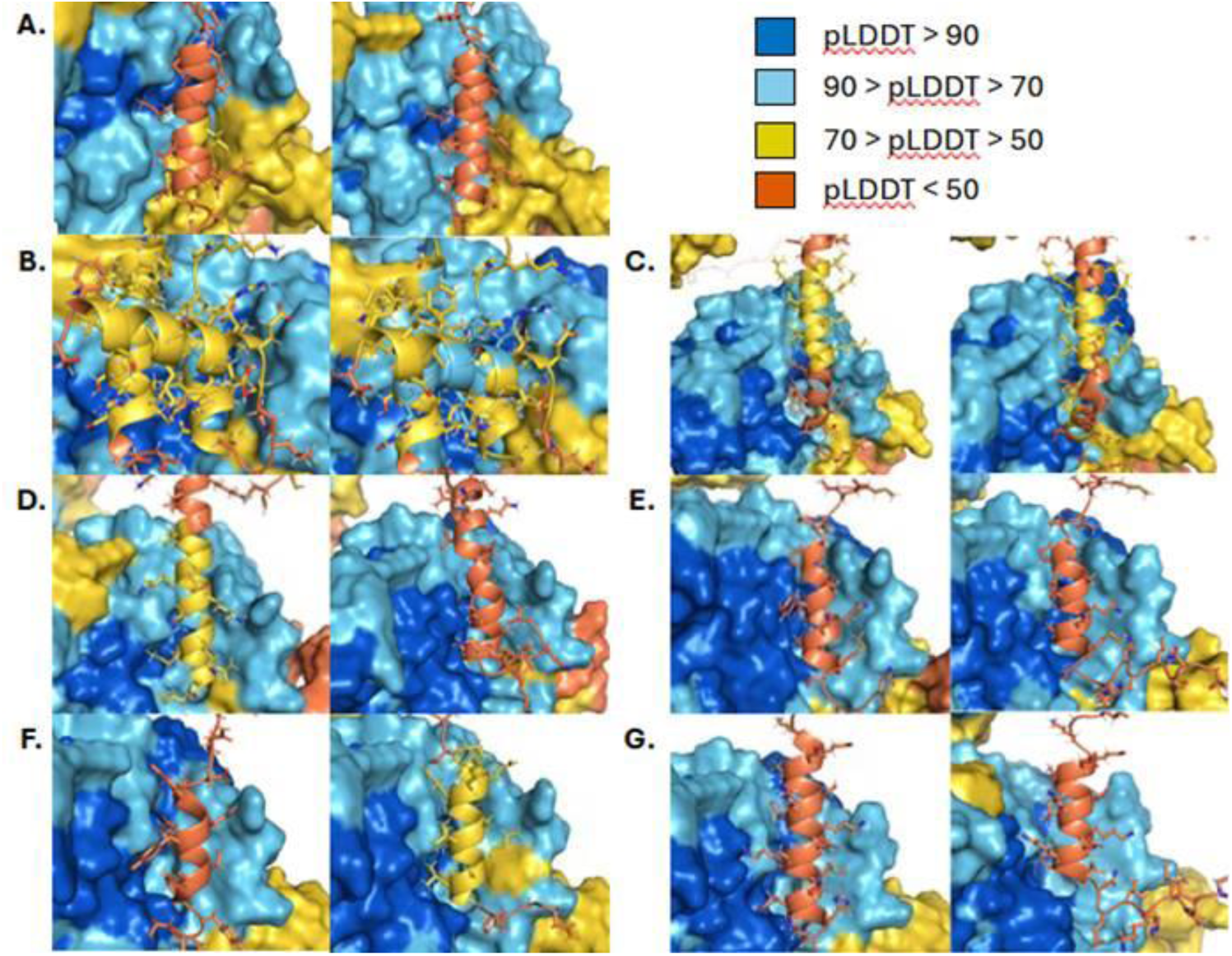
**pLDDT scores of the human proteins typically were unchanged or improved after truncations**. Shown here are the pLDDT scores of AlphaFold predictions of Spike S1, shown in surface view, and full-length (left) and truncated (right) human hits shown as cartoons with side chains. A) Paxillin; B) α-neoendorphin; C) BNIP3; D) α-synuclein; E) Presenilin-1; F) Protachykinin-1/Neuropeptide K; G) Synaptotagmin-1.

**Fig S4.**
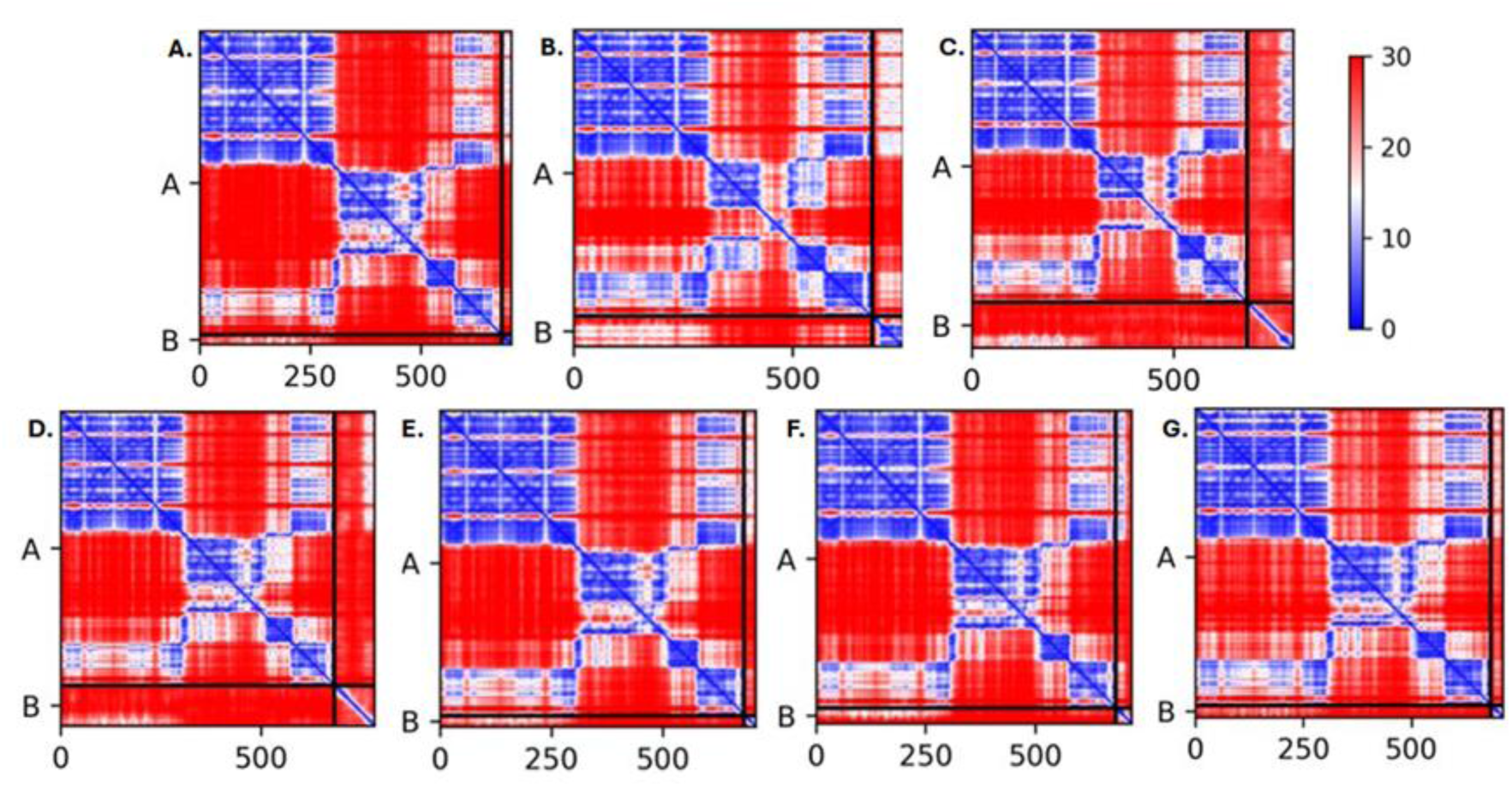
**Interacting regions of the human proteins and the spike NTD have much lower predicted aligned error**. AlphaFold output of predicted aligned error plots for Spike S1 and truncated human hits. A) Paxillin (Rank 5); B) α-neoendorphin (Rank 2); C) BNIP3 (Rank 3); D) α-synuclein (Rank 1); E) Presenilin-1 (Rank 1); F) Neuropeptide K (Rank 2); G) Synaptotagmin- 1 (Rank 1).

**Fig S5.**
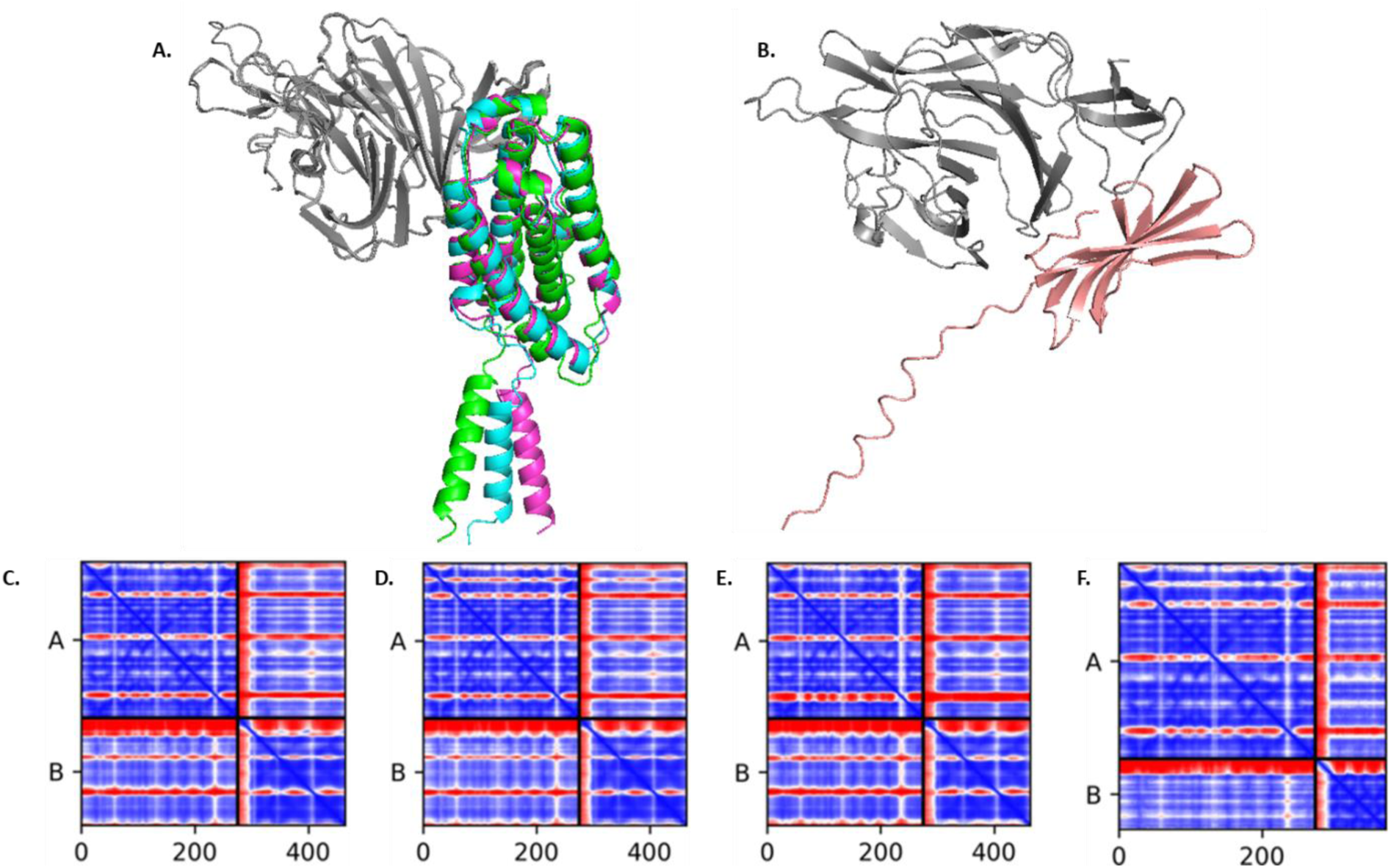
**Summit hits are predicted to bind near or close to the N-terminal domain’s hydrophobic cleft**. A) Superimposition of predicted complexes between NTD (grey) and interferon alpha-1/13 (P01562, blue), interferon alpha-14 (P01570, green), interferon alpha-2 (P01563, magenta); B) predicted complex between NTD (grey) and prostate and testis expressed protein 4 (P0C8F1, pink). Below are the predicted aligned error plots for the top five hits: C) P01562; D) P01570; E) P01563; F) P0C8F1.

**Fig S6.**
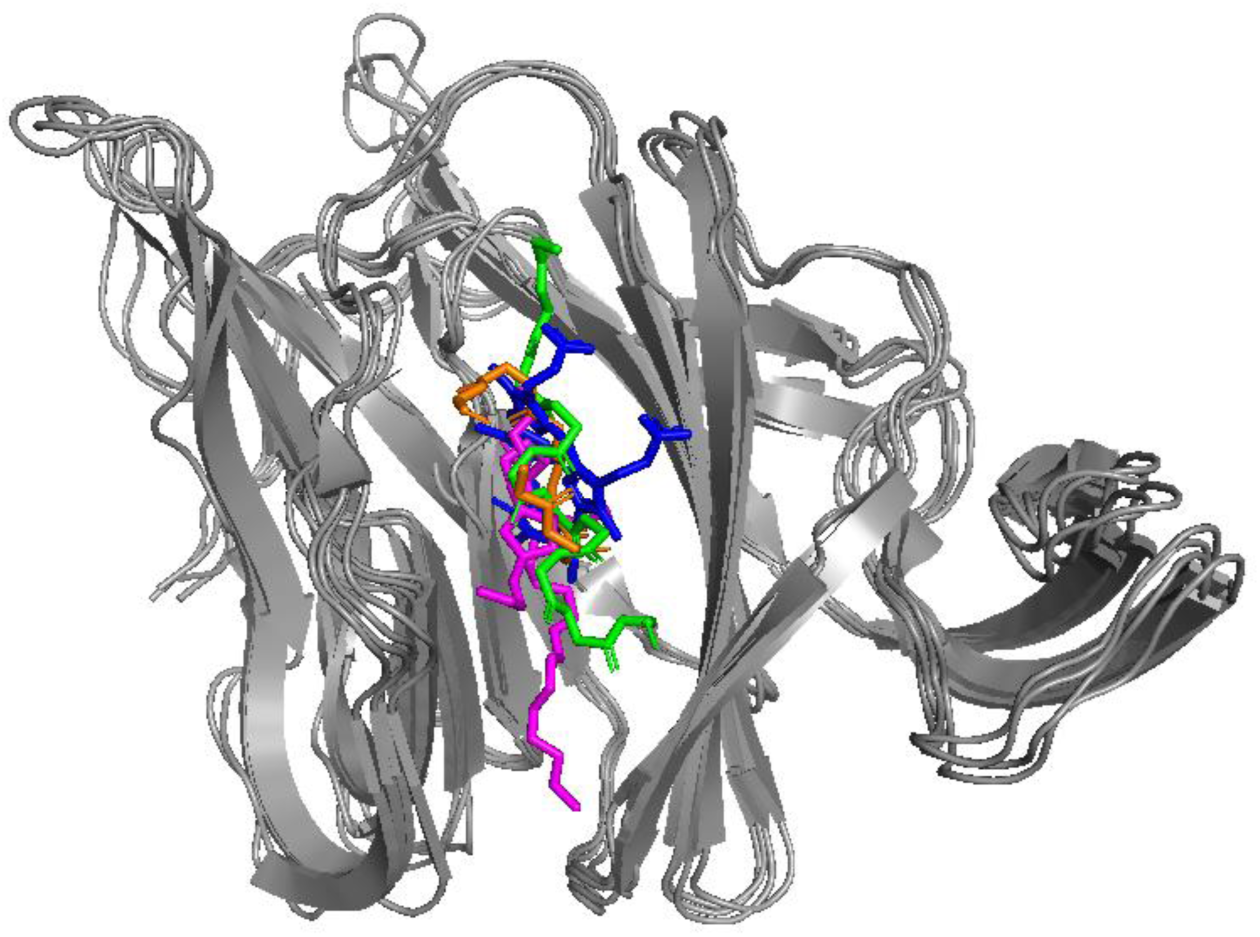
**Superimposition of multiple experimental NTD – ligand complexes suggest a common binding site of the hydrophobic cleft**. Heavy complementarity-determining region 3 loop of monoclonal antibody PVI.V6-14 Fab (7RBU, orange); Polysorbate 80 (7JJI, magenta); Biliverdin (7B62, blue); heavy complementarity-determining region H3 of neutralizing antibody 5-7 Fab (7RW2, green).

**Table S1.** Thirty-four PDBs released after the AlphaFold2 validation date were used to benchmark the workflow’s ability to predict viral – human protein complexes. . Here is the list of PDBs used in the benchmarking test set and the data output for the initial predictions. Asterisks in the RMSD column indicate misleading RMSD results arising from incorrect orientations, binding, and folding, no spatial proximity.

| PDB ID | Exp PISA (Å<sup>2</sup>) | Initial PISA (Å<sup>2</sup>) | Initial Clash | Initial pLDDT | Initial pTM | RMSD (Å) |
| --- | --- | --- | --- | --- | --- | --- |
| 6YXJ | 777.3 | 765.4 | 16.512 | 88.66 | 0.85 | 1.304 |
| 7CYV | 630.8 | 214.1 | 0 | 70.31 | 0.52 | ** |
| 7DDO | 941.5 | 1209.3 | 76.022 | 83.94 | 0.72 | ** |
| 7DHG | 1999.9 | 186.5 | 0.591 | 74.1 | 0.68 | ** |
| 7F15 | 561.8 | 211.7 | 0 | 76.07 | 0.53 | 23.646 |
| 7FG3 | 674.9 | 121.3 | 0 | 63.25 | 0.47 | 22.023 |
| 7LMS | 1220.1 | 996.7 | 48.255 | 81.91 | 0.68 | 1.064 |
| 7MIC | 2887.5 | 387.5 | 14.021 | 91.47 | 0.63 | 3.807 |
| 7NFR | 585.7 | 0 | 0 | 77.97 | 0.47 | 18.13 |
| 7P72 | 475.9 | 353.5 | 1.943 | 90.32 | 0.74 | ** |
| 7PKI | 885.5 | 992 | 79.855 | 85.63 | 0.73 | 16.919 |
| 7QTP | 397.4 | 154.5 | 0 | 83.93 | 0.72 | ** |
| 7RBR | 1022 | 197.1 | 0.647 | 90.19 | 0.75 | ** |
| 7RFY | 571.6 | 612.4 | 2.003 | 88.8 | 0.83 | ** |
| 7VMU | 1166.6 | 317.5 | 5.287 | 79.76 | 0.56 | 19.519 |
| 7VNB | 924.6 | 384.5 | 4.526 | 70.08 | 0.52 | 18.334 |
| 7WFC | 1178.9 | 199 | 4.159 | 77.28 | 0.65 | 20.783 |
| 7X6Y | 392.6 | 138.4 | 0 | 93.06 | 0.88 | ** |
| 7X6Z | 375.1 | 0 | 0 | 94.05 | 0.88 | ** |
| 7X70 | 377.1 | 0 | 0 | 94.47 | 0.89 | ** |
| 7XBH | 837 | 810.6 | 39.576 | 84.14 | 0.71 | ** |
| 7YA1 | 753.3 | 1187.5 | 74.747 | 86.03 | 0.73 | ** |
| 7YUE | 485.4 | 583.4 | 1.495 | 87.27 | 0.83 | 0.634 |
| 7ZCF | 1062.4 | 1744.2 | 119.39 | 58.85 | 0.4 | 16.78 |
| 7ZUD | 496.1 | 0 | 0 | 88.17 | 0.8 | ** |
| 8DCE | 789.6 | 338.2 | 14.155 | 72 | 0.54 | ** |
| 8DI5 | 951.2 | 778 | 43.779 | 60.03 | 0.42 | 18.261 |
| 8DLY | 999.7 | 50.8 | 0 | 63.48 | 0.46 | 19.457 |
| 8DM0 | 995.5 | 707.2 | 23.39 | 59.83 | 0.44 | 20.142 |
| 8F2P | 723.9 | 249.8 | 5.66 | 81.53 | 0.59 | ** |
| 8I4E | 1323.6 | 2327.1 | 101.417 | 60.08 | 0.41 | 12.748 |
| 8I4F | 1373.7 | 3577.3 | 227.787 | 57.36 | 0.39 | 17.9 |
| 8T12 | 1723.4 | 1607.4 | 71.732 | 79.68 | 0.55 | ** |
| 8T13 | 981.3 | 1607.4 | 71.732 | 79.68 | 0.55 | ** |

**Table S2.** Prediction statistics for the six selected hits. Displayed here are the hits (top) and negative controls (bottom). Asterisks in the RMSD column indicate misleading RMSD results arising from incorrect orientations, binding, and folding, no spatial proximity, etc. Dashes indicate no data.

| PDB ID | Rank of Model | Initial PISA (Å<sup>2</sup>) | Initial Clash | Initial pLDDT | Initial pTM | Initial RMSD (Å) | Relaxed PISA (Å<sup>2</sup>) | Relaxed Clash | Relaxed pLDDT | Relaxed pTM | Relaxed RMSD (Å) | MD PISA (Å<sup>2</sup>) | MD Clash | Relaxed RMSD (Å) |
| --- | --- | --- | --- | --- | --- | --- | --- | --- | --- | --- | --- | --- | --- | --- |
| 6YXJ | 1 | 765.4 | 16.512 | 88.66 | 0.85 | 1.304 | 716.5 | 0 | 89.68 | 0.87 | 1.558 | - | - | - |
|  | 2 | - | - | - | - | - | 720 | 0 | 89.06 | 0.87 | 1.314 | - | - | - |
|  | 3 | - | - | - | - | - | 699.2 | 0.995 | 88.76 | 0.87 | 1.773 | - | - | - |
|  | 4 | - | - | - | - | - | 771.4 | 0 | 88.62 | 0.85 | 1.28 | 959.2 | 0 | 1.964 |
|  | 5 | - | - | - | - | - | 749.5 | 0 | 88.39 | 0.85 | 1.279 | - | - | - |
| 7RFY | 1 | 612.4 | 2.003 | 88.8 | 0.83 | ** | 759 | 1.072 | 89.22 | 0.84 | ** | 881.2 | 0 | ** |
|  | 2 | - | - | - | - | - | 352.5 | 0 | 88.91 | 0.83 | ** | - | - | - |
|  | 3 | - | - | - | - | - | 576.3 | 0 | 88.88 | 0.83 | ** | - | - | - |
|  | 4 | - | - | - | - | - | 640.5 | 0 | 88.8 | 0.83 | ** | - | - | - |
|  | 5 | - | - | - | - | - | 548.9 | 0 | 88.51 | 0.82 | ** | - | - | - |
| 7YUE | 1 | 583.4 | 1.495 | 87.27 | 0.83 | 0.634 | 597.4 | 3.494 | 89.29 | 0.85 | 0.435 | 617.1 | 0 | 1.256 |
|  | 2 | - | - | - | - | - | 0 | 0 | 88.02 | 0.81 | ** | - | - | - |
|  | 3 | - | - | - | - | - | 584 | 0 | 87.27 | 0.83 | 0.62 | - | - | - |
|  | 4 | - | - | - | - | - | 576.7 | 0 | 87.16 | 0.74 | 0.336 | - | - | - |
|  | 5 | - | - | - | - | - | 824.8 | 4.797 | 84.26 | 0.71 | 1.479 | 624.7 | 0 | 3.513/** |
| 7XBH | 1 | 810.6 | 39.576 | 84.14 | 0.71 | ** | 551.2 | 2.417 | 88.67 | 0.73 | ** | - | - | - |
|  | 2 | - | - | - | - | - | 125 | 0 | 86.1 | 0.71 | ** | - | - | - |
|  | 3 | - | - | - | - | - | 14.9 | 0 | 85.13 | 0.73 | ** | - | - | - |
|  | 4 | - | - | - | - | - | 96.8 | 0.47 | 85 | 0.72 | ** | - | - | - |
|  | 5 | - | - | - | - | - | 797.2 | 2.163 | 84.12 | 0.71 | ** | 1627.8 | 0 | ** |
| 7LMS | 1 | 996.7 | 48.255 | 81.91 | 0.68 | 1.064 | 1500.8 | 0 | 82.33 | 0.81 | 0.986 | 1972.7 | 0.464 | 1.515 |
|  | 2 | - | - | - | - | - | 1004.8 | 4.527 | 81.9 | 0.68 | 1.076 | 1662.7 | 0 | ** |
|  | 3 | - | - | - | - | - | 988.2 | 1.971 | 80.49 | 0.68 | 1.141 | - | - | - |
|  | 4 | - | - | - | - | - | 388 | 0 | 80.33 | 0.68 | 1.024 | - | - | - |
|  | 5 | - | - | - | - | - | 1117.2 | 3.321 | 79.58 | 0.7 | 1.014 | - | - | - |
| 7X6Y | 1 | 138.4 | 0 | 93.06 | 0.88 | ** | 72.3 | 0 | 94.56 | 0.9 | ** | - | - | - |
|  | 2 | - | - | - | - | - | 71.6 | 0 | 94.12 | 0.89 | ** | - | - | - |
|  | 3 | - | - | - | - | - | 75.1 | 0 | 94.12 | 0.89 | ** | - | - | - |
|  | 4 | - | - | - | - | - | 0 | 0 | 93.74 | 0.88 | ** | - | - | - |
|  | 5 | - | - | - | - | - | 154.7 | 0 | 93.06 | 0.88 | ** | 314.4 | 0 | 0.731 |

**Table S3.** Analysis of variation in sequence length shows little impact on prediction accuracy. In some cases, however, a particular sequence length can help AlphaFold find the correct binding pose that other variations of the same sequence may miss, in this case the PDB (structural) sequence of 7DHG and the custom truncation of 7RFY. Asterisks indicate analysis calculated small PyMol RMSD, however visual inspection showed incorrect binding poses and dimeric structures.

| PDB ID | Exp PISA (Å<sup>2</sup>) | Initial PISA (Å<sup>2</sup>) | Initial RMSD (Å) | PDB Seq PISA | PDB Seq RMSD (Å) | Custom Trunc PISA | Custom Trunc RMSD (Å) | Uniprot Seq PISA | Uniprot Seq RMSD (Å) |
| --- | --- | --- | --- | --- | --- | --- | --- | --- | --- |
| 6YXJ | 777.3 | 765.4 | 1.304 | 643.9 | 1.839 | 328.4 | 0.463/0.469 |  |  |
| 7CYV | 630.8 | 214.1 | ** | 389.4 | ** | No Trunc | No Trunc | 378.4 | ** |
| 7DDO | 941.5 | 1209.3 | ** | 771.5 | ** | No Trunc | No Trunc | 798.2 | ** |
| 7DHG | 1999.9 | 186.5 | ** | 1067.2 | 2.137 | 1829.7 | **/** | 186.8 | ** |
| 7F15 | 561.8 | 211.7 | 23.646 | 86.5 | 23.882 | 0 | **/** | 988.5 | 28.076 |
| 7FG3 | 674.9 | 121.3 | 22.023 | 338.2 | 29.077 | 226.8 | 21.493/21.262 | 135.0 | 22.015 |
| 7LMS | 1220.1 | 996.7 | 1.064 | 1298 | 0.987 | 648.0 | **/** | 797.3 | ** |
| 7MIC | 2887.5 | 387.5 | 3.807 | 149.5 | 17.601 | No Trunc | No Trunc |  |  |
| 7NFR | 585.7 | 0 | 18.13 | 745.3 | 19.36 | 182.4 | 17.157/17.157 | 101.1 | 24.144 |
| 7P72 | 475.9 | 353.5 | ** | 312.5 | ** | 329.9 | 0.332/0.332 | 0? | 22.662 |
| 7PKI | 885.5 | 992 | 16.919 | 312.5 | 16.956 | 1294.2 | **/** | 73.9 | ** |
| 7QTP | 397.4 | 154.5 | ** | 421 | 0.476 | 0 | **/** | 261.2 | ** |
| 7RBR | 1022 | 197.1 | ** | 434.3 | ** | 607.5 | **/** | 428.4 | ** |
| 7RFY | 571.6 | 612.4 | ** | 586.2 | ** | 578.1 | 0.501/0.501 | 592.4 | 0.627 |
| 7VMU | 1166.6 | 317.5 | 19.519 | 244.6 | 24.086 | 156.9 | 22.583/22.539 | 1240.8 | 22.505 |
| 7VNB | 924.6 | 384.5 | 18.334 | 309.2 | 18.271 | 436.8 | 17.541/17.541 | 356.2 | 18.238 |
| 7WFC | 1178.9 | 199 | 20.783 | 309.2 | 20.919 | 126.8 | **/** | 480.3 | ** |
| 7X6Y | 392.6 | 138.4 | ** | 165 | ** | 232.7 | **/** | 50.1 | ** |
| 7X6Z | 375.1 | 0 | ** | 0 | ** | 318.3 | **/** | 89.7 | ** |
| 7X70 | 377.1 | 0 | ** | 0 | ** | 258.7 | **/** | 430.3 | ** |
| 7XBH | 837 | 810.6 | ** | 871.9 | ** | 612.4 | **/** | 317.7 | ** |
| 7YA1 | 753.3 | 1187.5 | ** | 871.9 | ** | 690.6 | **/** | 245.9 | 28.289 |
| 7YUE | 485.4 | 583.4 | 0.634 | 435.4 | ** | 59.6 | **/** | 192.5 | ** |
| 7ZCF | 1062.4 | 1744.2 | 16.78 | 392.8 | 23.049 | 842.5 | 20.622/20.622 | 1069.8 | 22.930 |
| 7ZUD | 496.1 | 0 | ** | 0 | ** | 0 | **/** | 1376.8 | ** |
| 8DCE | 789.6 | 338.2 | ** | 426.8 | ** | 19.6 | **/** | 2860 | ** |
| 8DI5 | 951.2 | 778 | 18.261 | 636.8 | 11.045 | 759.6 | **/** | 420.3 | 13.418 |
| 8DLY | 999.7 | 50.8 | 19.457 | 509.7 | 10.600 | 565.2 | **/** | 55.6 | 18.588 |
| 8DM0 | 995.5 | 707.2 | 20.142 | 413.8 | 10.651 | 701.1 | **/** | 58.8 | 18.080 |
| 8F2P | 723.9 | 249.8 | ** | 112.5 | ** | 218.6 | 8.779/8776 | 610.9 | 21.442 |
| 8I4E | 1323.6 | 2327.1 | 12.748 | 615.6 | 18.59 | 465.1 | 30.822/21.992 | 1493.9 | 13.431 |
| 8I4F | 1373.7 | 3577.3 | 17.9 | 113.7 | ** | 594.2 | 30.732/20.689 | 918.3 | 12.754 |
| 8T12 | 1723.4 | 1607.4 | ** | 1500.5 | ** | 65.5 | **/** | 3151.8 | ** |
| 8T13 | 981.3 | 1607.4 | ** | 408.8 | ** | 116.8 | **/** | 1609.6 | ** |

**Table S4.** Values of the filtering criteria for the selected models typically improved throughout the workflow. Displayed is all numerical data of the models during the course of the workflow when applied to the small-scale cell junction and synaptic protein screen. Dashes indicate no data.

| Protein | Uniprot ID | Rank | Initial PISA (Å<sup>2</sup>) | Initial Clash | Relaxed PISA (Å<sup>2</sup>) | Relaxed Clash | Truncated PISA (Å<sup>2</sup>) | Truncated Clash | Final pLDDT | Triplicate MD PISA (Å<sup>2</sup>) | Triplicate MD Clash |
| --- | --- | --- | --- | --- | --- | --- | --- | --- | --- | --- | --- |
| Paxillin | A0A1B0GV30 | Rank 1 | 748.2 | 17.54 | 314.7 | 0.92 | 486.3 | 2.03 | 74.49 | - | - |
|  |  | Rank 2 | - | - | 145.4 | 0 | 232.6 | 0 | 74.25 | - | - |
|  |  | Rank 3 | - | - | 619.4 | 8.57 | 496.7 | 1.63 | 71.96 | - | - |
|  |  | Rank 4 | - | - | 404.6 | 0 | 225.5 | 0 | 70.99 | - | - |
|  |  | Rank 5 | - | - | 673 | 8.93 | 590.2 | 2.22 | 68.46 | 987.9 | 0 |
| BNIP3 | Q12983 | Rank 1 | 941.6 | 16.36 | 1141.8 | 46.79 | 898.3 | 2.79 | 68.62 | - | - |
|  |  | Rank 2 | - | - | 1054.6 | 13.58 | 893.1 | 0.86 | 67.62 | - | - |
|  |  | Rank 3 | - | - | 831.8 | 0 | 947.9 | 0.91 | 67.59 | 1181.5 | 0 |
|  |  | Rank 4 | - | - | 739.3 | 72.84 | 907.4 | 2.69 | 66.36 | - | - |
|  |  | Rank 5 | - | - | 751.8 | 17.71 | 895.2 | 0.45 | 66.22 | - | - |
| Protachykinin-1 | P20366 | Rank 1 | 912.5 | 15.49 | 1163.5 | 25.26 | 752.3 | 1.53 | 73.11 | - | - |
|  |  | Rank 2 | - | - | 1075.2 | 42.14 | 772 | 0.54 | 72.19 | 938.7 | 0 |
|  |  | Rank 3 | - | - | 1478.5 | 60.03 | 673.8 | 0.59 | 71.82 | - | - |
|  |  | Rank 4 | - | - | 674.6 | 17.03 | 207.9 | 0 | 70.06 | - | - |
|  |  | Rank 5 | - | - | 38.8 | 0 | 480.6 | 0 | 69.25 | - | - |
| Alpha-neoendorphin | A0A494C0B1 | Rank 1 | 699.6 | 13.77 | 405.1 | 3.58 | 139.1 | 0 | 75.06 | - | - |
|  |  | Rank 2 | - | - | 798.4 | 17.94 | 834.4 | 0.48 | 71.19 | 851.7 | 0.47 |
|  |  | Rank 3 | - | - | 56.4 | 0 | 91.6 | 0 | 70.51 | - | - |
|  |  | Rank 4 | - | - | 700.7 | 13.61 | 29.2 | 0 | 69.94 | - | - |
|  |  | Rank 5 | - | - | 766.3 | 11.46 | 354.2 | 0 | 66.54 | - | - |
| Alpha-synuclein | A0A669KB41 | Rank 1 | 1000.9 | 16.49 | 699.8 | 28.39 | 610.1 | 0.92 | 68.38 | 1246.8 | 0 |
|  |  | Rank 2 | - | - | 774.6 | 24.88 | 180.4 | 0 | 66.82 | - | - |
|  |  | Rank 3 | - | - | 0 | 0 | 0 | 0 | 66.79 | - | - |
|  |  | Rank 4 | - | - | 453.9 | 33.41 | 556.6 | 1.99 | 65.48 | - | - |
|  |  | Rank 5 | - | - | 31.5 | 0 | 261.4 | 0 | 62.38 | - | - |
| Synaptotagmin-1 | F8VYH8 | Rank 1 | 749.3 | 14.54 | 648.4 | 10.59 | 459.1 | 1.08 | 73.63 | 1082.4 | 0 |
|  |  | Rank 2 | - | - | 361.2 | 4.84 | 140.3 | 0 | 72.93 | - | - |
|  |  | Rank 3 | - | - | 564.6 | 4.33 | 232.4 | 0 | 72.79 | - | - |
|  |  | Rank 4 | - | - | 323.9 | 6.68 | 279.8 | 0 | 70.95 | - | - |
|  |  | Rank 5 | - | - | 646.9 | 10.52 | 134.6 | 0 | 70.61 | - | - |
| Presenilin-1 | G3V3P0 | Rank 1 | 610 | 12.20 | 712.9 | 19.54 | 774.3 | 0.67 | 73.41 | 1150.9 | 0 |
|  |  | Rank 2 | - | - | 319.5 | 0 | 719.9 | 0.40 | 70.39 | - | - |
|  |  | Rank 3 | - | - | 449 | 3.76 | 760.8 | 1.81 | 70.08 | - | - |
|  |  | Rank 4 | - | - | 675.9 | 13.82 | 326.2 | 0 | 69.79 | - | - |
|  |  | Rank 5 | - | - | 458.3 | 6.12 | 464.3 | 1.02 | 68.53 | - | - |

**Table S5.** Seventeen hits were found during the proteome screening. . Displayed are the numerical data of the best models of the resulting hits when applying the workflow to the human proteome.

| Name | Uniprot ID | Best Visual Model | PISA (Å<sup>2</sup>) | Total Clash | Avg Clash | NTD pLDDT | Ligand pLDDT | Total pLDDT |
| --- | --- | --- | --- | --- | --- | --- | --- | --- |
| Interferon alpha-1/13 | P01562 | rank_1_model_3 | 885.3 | 0.886 | 0.443 | 85.97 | 81.15 | 83.96 |
| Interferon alpha-14 | P01570 | rank_1_model_3 | 862 | 0.432 | 0.432 | 84.52 | 81.22 | 83.12 |
| Interferon alpha-2 | P01563 | rank_1_model_3 | 963.9 | 0.411 | 0.411 | 83.53 | 80.58 | 82.29 |
| Leukocyte immunoglobulin-like receptor subfamily A member 6 | Q6PI73 | rank_2_model_5 | 530.7 | 0 | 0 | 80.49 | 81.47 | 81.09 |
| Prostate and testis expressed protein 3 | B3GLJ2 | rank_3_model_4 | 608.9 | 0 | 0 | 81.82 | 82.19 | 81.88 |
| Prostate and testis expressed protein 4 | P0C8F1 | rank_1_model_4 | 972.8 | 0.819 | 0.409 | 85.03 | 80.65 | 83.83 |
| PRUNE1 | Q86TP1 | rank_3_model_1 | 861.4 | 3.472 | 0.496 | 80.85 | 81.09 | 80.98 |
| Sorting nexin-6 | Q9UNH7 | rank_1_model_5 | 963.5 | 1.072 | 0.536 | 82.11 | 84.19 | 83.33 |
| T cell receptor alpha variable 14/delta variable 4 | A0A0A6YYC5 | rank_2_model_3 | 600.2 | 1.633 | 0.544 | 85.06 | 80.85 | 83.79 |
| T cell receptor alpha variable 38-1 | A0A0B4J264 | rank_1_model_4 | 607.4 | 0.567 | 0.567 | 85.52 | 81.11 | 84.17 |
| T cell receptor beta variable 12-5 | A0A1B0GX78 | rank_2_model_5 | 522 | 1.267 | 0.634 | 83.37 | 80.14 | 82.39 |
| T cell receptor beta variable 24-1 | A0A075B6N3 | rank_5_model_1 | 527.4 | 0 | 0 | 81.60 | 83.71 | 82.18 |
| T cell receptor beta variable 3-1 | A0A576 | rank_1_model_4 | 543.4 | 0.464 | 0.464 | 86.17 | 85.01 | 85.79 |
| Tetratricopeptide repeat protein 39A | Q5SRH9 | rank_2_model_5 | 632.1 | 0.427 | 0.427 | 81.37 | 83.28 | 82.67 |
| Translocon-associated protein subunit delta | P51571 | rank_1_model_5 | 542.9 | 1.3 | 0.433 | 84.52 | 83.40 | 84.06 |
| Trefoil factor 2 | Q03403 | rank_3_model_3 | 738.9 | 1.023 | 0.512 | 82.85 | 83.13 | 82.89 |

